# Dual modes of DNA N^6^-methyladenine maintenance by distinct methyltransferase complexes

**DOI:** 10.1101/2024.07.21.604504

**Authors:** Yuanyuan Wang, Bei Nan, Fei Ye, Zhe Zhang, Wentao Yang, Bo Pan, Junhua Niu, Aili Ju, Yongqiang Liu, Wenxin Zhang, Yifan Liu, Shan Gao

**Author notes:** Corresponding author. Yuanyuan Wang, Yifan Liu; Shan Gao. These authors contributed equally (Yuanyuan Wang, Bei Nan, Fei Ye, Zhe Zhang, Wentao Yang).

## Abstract

Stable inheritance of DNA N^6^-methyladenine (6mA) is crucial for its biological functions in eukaryotes. Here, we identify two distinct methyltransferase (MTase) complexes, both sharing the catalytic subunit AMT1, but featuring AMT6 and AMT7 as their unique components, respectively. While the two complexes are jointly responsible for 6mA maintenance methylation, they exhibit distinct enzymology, DNA/chromatin affinity, genomic distribution, and knockout phenotypes. AMT7 complex, featuring high MTase activity and processivity, is connected to transcription-associated epigenetic marks, including H2A.Z and H3K4me3, and is required for the bulk of maintenance methylation. In contrast, AMT6 complex, with reduced activity and processivity, is recruited to initiate maintenance methylation immediately after DNA replication. These two complexes coordinate in maintenance methylation. By integrating signals from both replication and transcription, this mechanism ensures the faithful and efficient transmission of 6mA as an epigenetic mark in eukaryotes.

**Significance statement:** DNA N^6^-methyladenine (6mA) has recently been recognized as an epigenetic mark in eukaryotes. The stable inheritance of 6mA is essential for its biological functions. However, the precise mechanisms by which 6mA patterns are faithfully and efficiently transmitted remain largely unknown. Here, we have identified two distinct 6mA methyltransferase (MTase) complexes and elucidated their coordinated role in maintenance methylation. This dual- complex mechanism ensures rapid and accurate methylation at newly replicated loci with proper transcription-associated epigenetic marks.

## Introduction

DNA N^6^-methyladenine (6mA) is well-documented in prokaryotes, serving crucial functions in maintaining genome integrity and regulating replication and transcription (1). Recent studies have unveiled its pervasive presence in eukaryotes, albeit with notable variation in abundance and functionality across species (2–5). In animals, plants, and higher fungi, 6mA is sparse, distributed with no specific sequence context, and frequently associated with transcriptionally inactive regions, raising questions over whether it serves as an enzymatically deposited epigenetic mark (6–15). By contrast, in protists, green algae, and basal fungi, 6mA is abundant, present preferentially, if not exclusively, at the ApT dinucleotide, and associated with RNA polymerase II (Pol II)-transcribed genes, supporting its role as an epigenetic mark (3–5, 16–18).

As an epigenetic mark, 6mA must be stably propagated. The inheritance mechanism of 6mA is a crucial aspect of 6mA biology in eukaryotes. We and others have recently established 6mA as an enzymatically deposited and heritable epigenetic modification in the unicellular eukaryote *Tetrahymena thermophila* (16, 19–21). 6mA is faithfully transmitted through a semiconservative mechanism dependent on an MT-A70 family member, AMT1 (adenine methyltransferase 1) (19). It is enriched towards the 5’ end of the gene body (21) and on linker DNA flanked by nucleosomes decorated with the histone variant H2A.Z and the histone modification H3 lysine 4 trimethylation (H3K4me3) (20). Reduced 6mA level alters gene expression, diminishes growth, and generates defects in morphology and behavior (20). However, how this stereotypical 6mA distribution pattern is faithfully and efficiently transmitted remains unclear. The connection of 6mA with other epigenetic marks, and the coordination between the methylation process on the one hand, and DNA replication and transcription on the other, still need to be explored.

Here we characterize two AMT1-containing methyltransferase (MTase) complexes, distinguished by the inclusion of two mutually exclusive components, AMT6 and AMT7. AMT7 complex exhibits high MTase activity and processivity, with its distribution strongly correlated with transcription- associated epigenetic marks, especially H2A.Z and H3K4me3. It is recruited by these epigenetic pathways and required for most of maintenance methylation. In contrast, AMT6 complex exhibits reduced MTase activity and processivity, and is more widely distributed. It is recruited to facilitate the initiation of maintenance methylation after DNA replication. This dual-mode system achieves precise and rapid methylation through multiple recognitions, integrates signals from both replication and transcription, and ensures transmission of 6mA as an epigenetic mark.

## Results

### Two distinct AMT1 complexes with AMT6 or AMT7 as their mutually exclusive components

We performed immunoprecipitation and mass spectrometry (IP-MS) using AMT1 as bait (Fig. S1A). In addition to the previously reported interacting proteins, namely AMT7, AMTP1, and AMTP2 (referred to as MTA9, p1, and p2 in (16)), we also identified AMT6 (referred to as MTA9-B in (16)), another MT- A70 family member (Fig. 1A). We then performed reciprocal IP with AMT6 and AMT7 as bait. MS revealed that both AMT6 and AMT7 interacted with AMT1, AMTP1, and AMTP2 (Fig. 1A). These interactions were corroborated by immunoblotting (Fig. 1B). Importantly, in AMT7 IP, we could not detect AMT6, and *vice versa* (Fig. 1A, C). Therefore, even though AMT6 and AMT7 share their core interactomes, they form two distinct and mutually exclusive complexes: AMT6-AMT1-AMTP1-AMTP2 (hereafter referred to as AMT6 complex) and AMT7-AMT1-AMTP1-AMTP2 (AMT7 complex) (Fig. 1D).

**Fig. 1.**
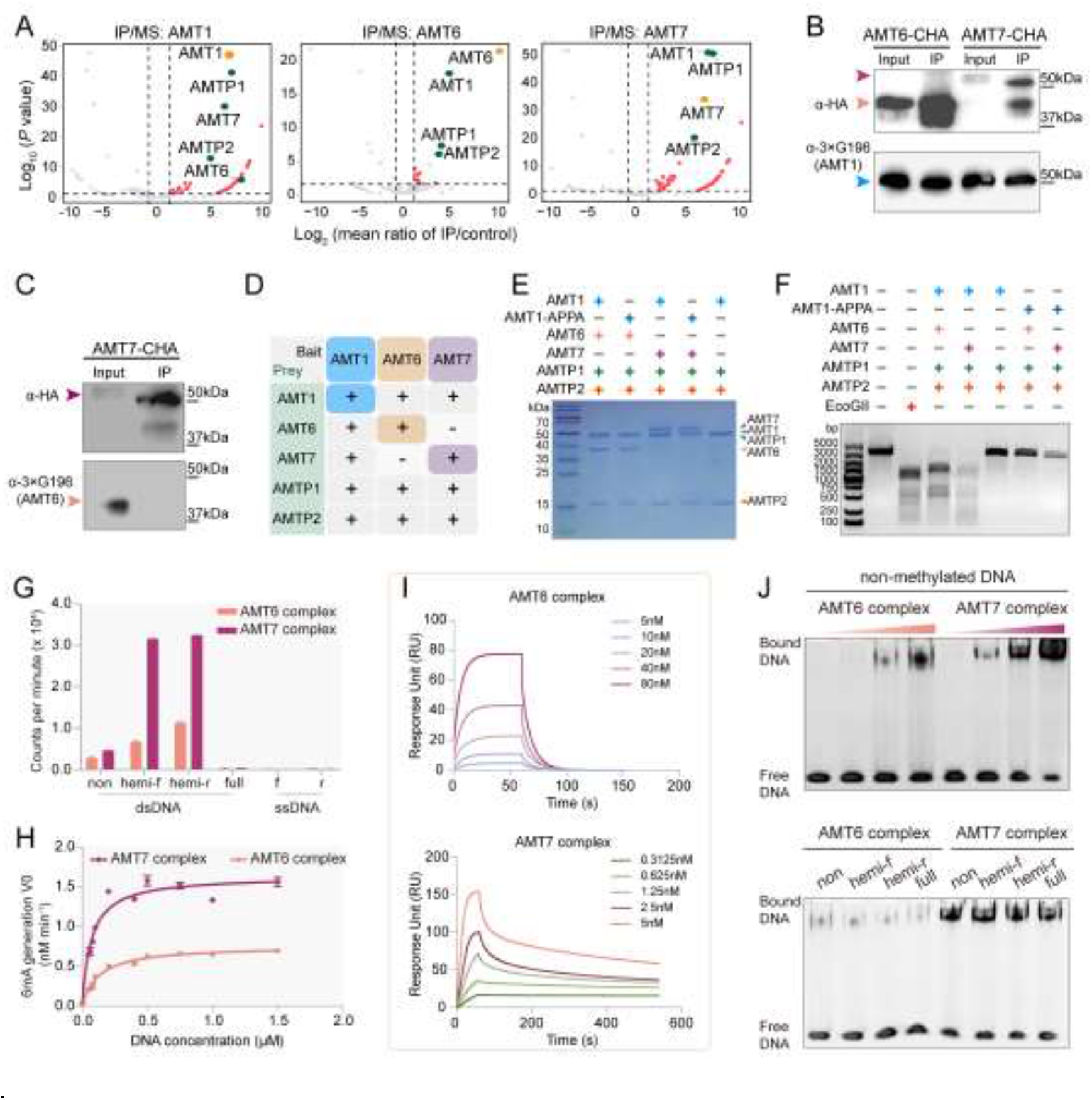
Two distinct AMT1 complexes with AMT6 or AMT7 as their mutually exclusive components. A. Volcano plots of IP-MS results using AMT1, AMT6, and AMT7 as the bait. Bait: orange; high-confidence preys of interest: green; other high- confidence preys: red; low-confidence preys: gray. B. IP-IB confirming the interaction of AMT1 with AMT6 and AMT7, respectively. Bait proteins were tagged with a hemagglutinin (HA), while prey proteins were tagged with 3×G196. C. IP-IB confirming the lack of interaction between AMT6 and AMT7. Bait proteins were HA-tagged, while prey proteins were tagged with 3×G196. D. Summary of interactions between AMT1 complexes according to IP-MS results. E. SDS-PAGE analysis of recombinant AMT6 and AMT7 complexes with bacterially expressed proteins. F. Detection of DNA methyltransferase (MTase) activities by *Dpn*I digestion. Linearized plasmid DNA purified from *dam- E. coli* was subject to *in vitro* methylation. Only methylated DNA was digested by *Dpn*I, as seen in lanes treated by wild-type AMT6 and AMT7 complexes, as well as M.EcoGII. G. Detection of DNA MTase activities by [^3^H]SAM labeling. 27bp double- strand DNA (dsDNA) substrates, with a single ApT dinucleotide in the non-methylated state (non), hemi-methylated on the forward strand (hemi-f), hemi-methylated on the reverse strand (hemi-r), or fully methylated (full), were subject to *in vitro* methylation by AMT6 and AMT7 complexes. DNA methylation was monitored by scintillation counting. Only background methylation was observed in fully-methylated dsDNA, or single-strand DNA (ssDNA: f/r). Error bars represent standard deviation (S.D.) (n = 3). H. Enzyme kinetics of AMT6 and AMT7 complexes on hemi-methylated DNA, quantified by [^3^H]SAM-labeling. Each data point was presented as mean ± S.D. (n=3). Curve fitting was based on steady-state Michaelis- Menten kinetics. I. Detection of DNA binding by Surface Plasmon Resonance (SPR). Hemi- methylated dsDNA was immobilized, while specified concentrations of AMT6 and AMT7 complexes were flowed. Equilibrium and kinetic constants were calculated by fitting to 1:1 Langmuir binding model. J. Detection of DNA binding by Electrophoretic Mobility Shift Assay (EMSA). Top panel: bound DNA (top band) progressively increased, while free DNA (bottom) progressively decreased, as AMT6 and AMT7 complexes increased in concentration (1, 2, 4, 8 μM protein, 4 μM DNA); DNA binding ability was more prominent for AMT7 complex than AMT6 complex. 27bp non-methylated DNA was used as substrate. Bottom panel: no substantial difference between dsDNA of different methylation states.

AMT7 complex corresponds to the originally identified and characterized AMT1 complex (16, 19). However, AMT6 has been reported to fractionate separately from AMT1 (16), thus requiring further scrutiny. *AMT6* expression was elevated during vegetative growth (asexual reproduction) and conjugation (sexual reproduction), but diminished during starvation (cell cycle arrest), in a pattern very similar to that of AMT7, as well as AMT1, AMTP1, and AMTP2 (Fig. S1B). Like AMT1 and AMT7, AMT6 was also localized in the transcriptionally active macronucleus (MAC) but not in the transcriptionally inert micronucleus (MIC) during vegetative growth, as shown by immunofluorescence (IF) staining (Fig. S1C). During conjugation, AMT6 was detected in the parental MAC at pre- zygotic stages and then in the new MAC at post-zygotic stages (Fig. S1D). This localization pattern is consistent with that of 6mA (20, 22), supporting AMT6’s role in 6mA transmission.

Phylogenetic analysis grouped AMT6 and AMT7 in the same clade of the MT-A70 family, separate from the AMT1 clade (Fig. S1E) (20). Both AMT6 and AMT7 lacked the conserved MTase signature motif ([DNSH]PP[YFW]) required for catalytic activity (NIKW in AMT6 and NALW in AMT7) (Fig. S1F) (23), which is present in AMT1 (DPPW). AMT6 and AMT7 homologues were found in a wide range of unicellular eukaryotes, but absent in multicellular eukaryotes, mirroring the phylogenetic distribution of AMT1 (Fig. S1E) (20).

### Differential activities of reconstituted AMT6 and AMT7 complexes

We next reconstituted AMT6 and AMT7 complexes with bacterially expressed proteins (Fig. 1E). Their components co-fractionated during size exclusion chromatography (Fig. S1G), consistent with the formation of two stable complexes. Both complexes showed robust MTase activities, assayed by *Dpn*I digestion of methylated plasmid DNA (Fig. 1F) or scintillator count after ^3^H-SAM labeling of short double-stranded DNA (dsDNA) (Fig. 1G). For both complexes, we also observed higher MTase activities on the hemi-methylated substrate compared to the non-methylated substrate (Fig. 1G), in line with their role in maintenance methylation (19). No activity was observed for single-stranded DNA (ssDNA) (Fig. 1G). The MTase activity for AMT7 complex was substantially higher than AMT6 complex, as quantified by the steady-state kinetics using the hemi-methylated short dsDNA substrate (AMT7: Km=0.069μM, kcat=0.033min^-1^; AMT6: Km=0.126μM, kcat=0.015min^-1^) (Fig. 1H).

Notably, AMT1, AMTP1, and AMTP2 formed a stable subcomplex (hereafter referred to as AMT1 subcomplex; Fig. 1E). However, in the absence of AMT6 and AMT7, it did not have any MTase activity; neither did an AMT7 complex with AMT1 mutated at the MTase signature motif (DPPW to APPA) (Fig. 1F). Therefore, even though AMT1 subcomplex contains the catalytically critical subunit, it only becomes active after forming a full complex by incorporating either AMT6 or AMT7.

The difference in MTase activities of AMT6 and AMT7 complexes can be at least partially attributed to their different DNA binding affinity. Binding was specific for dsDNA (Fig. S1H), regardless of its methylation state (Fig. 1J, bottom panel). AMT7 complex showed a much higher DNA binding affinity than AMT6 complex, as revealed by Surface Plasmon Resonance (SPR) (Fig. 1I) and Electrophoretic Mobility Shift Assay (EMSA) (Fig. 1J). The disassociation constant (KD), as quantified by SPR, was much lower for AMT7 complex (KD=2.315×10^-10^M; ka=6.688×10^6^/Ms and kd=1.548×10^-3^/s) than for AMT6 complex (KD=6.498×10^-6^M; ka=3.894×10^4^/Ms and kd=0.2531/s) (Fig. 1I). Notably, EMSA showed that AMT1 subcomplex had low DNA binding affinity (Fig. S1I). Also, AMT7 alone did not bind DNA as tightly as AMT7 complex (Fig. S1J). Recent structural analyses of AMT7 complex support that AMT1 serves as the catalytic subunit, while AMT7 engages the substrate DNA from the opposite side (24, 25). Building on this structural insight and our results, we propose that the clamp-like structure formed by AMT7 complex wrapping around the dsDNA substrate may be more stable than its counterpart formed by AMT6 complex, leading to their differential DNA binding affinity, MTase activity, and processivity (see **Distinct 6mA profiles of Δ*AMT6* and Δ*AMT7***).

### Divergent phenotypes of Δ*AMT6* and Δ*AMT7*

Having identified AMT6 and AMT7 complexes and characterized their differential activities, we next performed loss-of-function genetic studies of AMT6 and AMT7. Δ*AMT7* largely phenocopied Δ*AMT1* (20): along with substantial global reduction of 6mA, as quantified by IF and MS (Fig. S2A, B), these mutants grew much more slowly than WT (2.6× WT doubling time) (Fig. S2C); many cells contained a dysfunctional contractile vacuole (Fig. 2A). In contrast, Δ*AMT6* behaved more like WT, with global 6mA level, growth rate, and contractile vacuole barely affected (Fig. 2A, Fig. S2A-C).

**Fig. 2.**
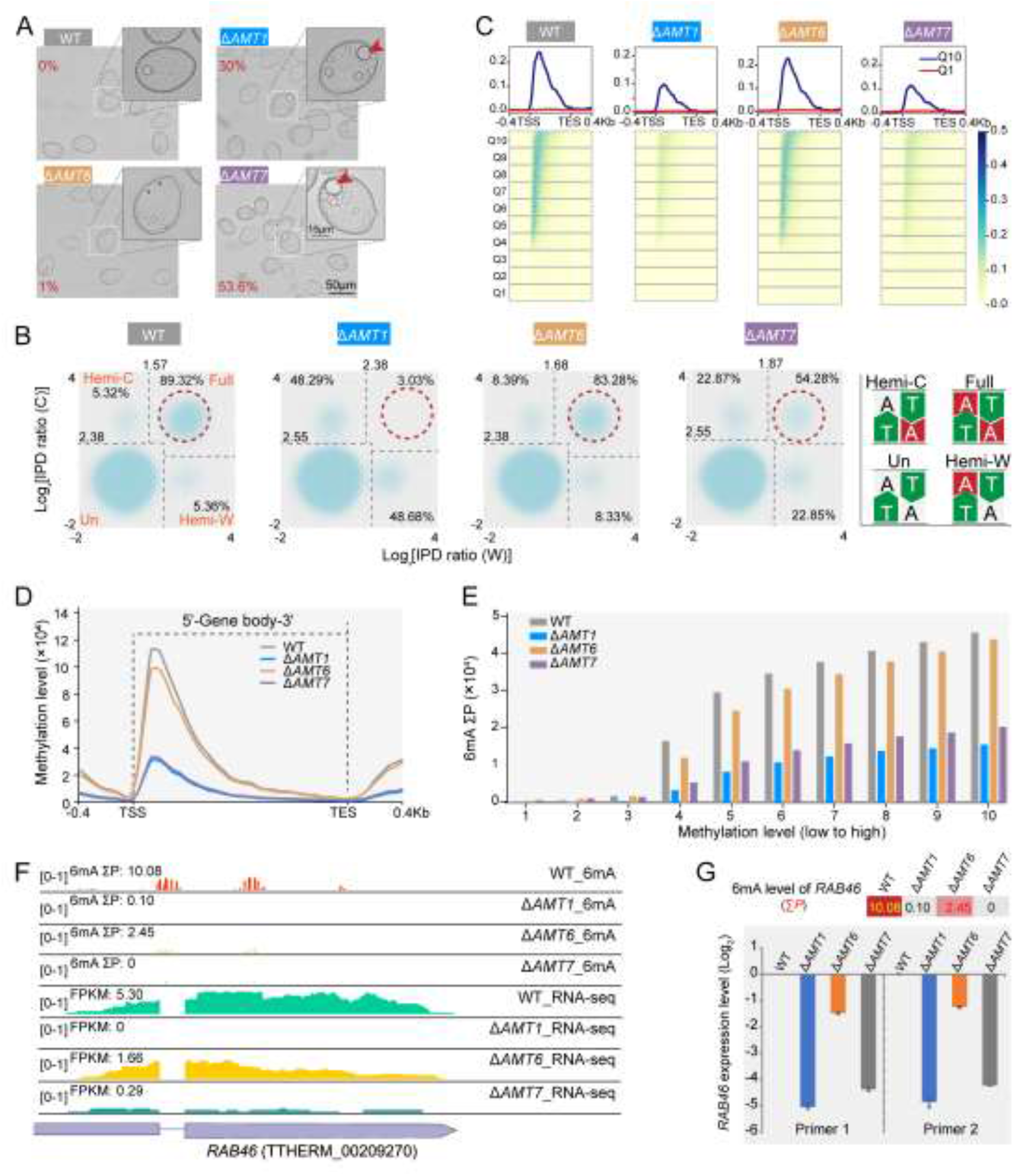
Distinct phenotypes of Δ*AMT6* and Δ*AMT7*. A. Representative bright-field images of WT, Δ*AMT1*, Δ*AMT6*, and Δ*AMT7*. Inset: zoom-in showing abnormally large contractile vacuoles (CV) frequently observed in Δ*AMT1* and Δ*AMT7*(red arrowheads), but not in WT and Δ*AMT6*. Percentage of abnormally large CV-containing cells was marked. B. Full-6mApT and hemi-6mApT levels in WT, Δ*AMT1*, Δ*AMT6*, and Δ*AMT7*. Note the abundance of full methylation in WT and Δ*AMT6*, but its absence in Δ*AMT1* and substantial reduction in Δ*AMT7*. C. 6mA distribution along Pol II-transcribed genes in WT, Δ*AMT1*, Δ*AMT6*, and Δ*AMT7*. Genes were ranked from low to high (quantiles 1-10, Q1-Q10) according to their methylation levels (ΣP, sum of penetrance values for all methylated ApT positions in the gene body) in WT. Genes were scaled to unit length and extended to each side for 0.4Kb. TSS, transcription start site. TES, transcription end site. D. 6mA distribution on the gene body of Pol II-transcribed genes in WT, Δ*AMT1*, Δ*AMT6*, and Δ*AMT7*. E. 6mA methylation level of 10 quantiles in WT, Δ*AMT1*, Δ*AMT6*, and Δ*AMT7*. Y-axis: ΣP for all methylated ApT positions in the gene body of a specified quantile. X-axis: 10 quantiles of genes ranked by their 6mA methylation level (1-10, from low to high). F. 6mA (SMRT CCS) and expression levels (RNA-seq) of *RAB46* in WT, Δ*AMT1*, Δ*AMT6*, and Δ*AMT7*. G. 6mA (ΣP) and expression levels (qRT-PCR) of *RAB46* in WT, Δ*AMT1*, Δ*AMT6*, and Δ*AMT7*.

Using Single Molecule, Real-Time (SMRT) sequencing in the Circular Consensus Sequences (CCS) mode (PacBio HiFi sequencing), we analyzed native genomic DNA from vegetatively growing Δ*AMT6* and Δ*AMT7* cells (1,259,892 and 730,284 high-quality reads, corresponding to 31× and 27× coverage of the *Tetrahymena* MAC genome, respectively). Essentially all 6mA occurred on the ApT dinucleotide (6mApT) (Fig. S2D, E), consistent with previously reported observations in WT and Δ*AMT1* (19). Importantly, we were able to distinguish between full-6mApT and hemi-6mApT (Fig. 2B, Fig. S2F, Table S1). Full-6mApT was predominant over hemi-6mApT in WT (full: 89.32%; hemi: 10.68%), but barely present in Δ*AMT1* (full: 3.03%; hemi: 96.97%) (Fig. 2B, Table S1). The former reflects complete and rapid maintenance methylation, while the latter is the result of *de novo* methylation. In Δ*AMT7*, full-6mApT and hemi-6mApT were at similar levels (full: 54.28%; hemi: 45.72%), reflecting incomplete maintenance methylation (Fig. 2B, Table S1). A more moderate increase in hemi-methylation was observed in Δ*AMT6*, with a corresponding decrease in full-methylation (full: 83.28%; hemi: 16.72%), reflecting delayed maintenance methylation (see **Distinct 6mA profiles of Δ*AMT6* and Δ*AMT7***) (Fig. 2B, Table S1).

We next examined 6mA distribution along Pol II-transcribed genes. In Δ*AMT6* and Δ*AMT7*, 6mA was still enriched towards the 5’ end of the gene body (Fig. 2C, D, Fig. S2G). 6mA levels across the gene body were slightly lower in Δ*AMT6* compared to WT; 6mA levels in Δ*AMT7* were much lower than WT, but slightly higher than Δ*AMT1* (Fig. 2C, D). High-6mA genes in WT showed reduction in both Δ*AMT6* and Δ*AMT7*; the decrease was often much more severe in Δ*AMT7* than in Δ*AMT6* (Fig. 2C, E). Notably, *RAB46*, encoding a Rab family GTPase regulating membrane fusion/fission (26, 27), contained substantial 6mA levels in WT and Δ*AMT6*, but lost almost all 6mA in Δ*AMT1* and Δ*AMT7* (Fig. 2F, G). *RAB46* expression levels were also greatly diminished in Δ*AMT1* and Δ*AMT7*, relative to WT and Δ*AMT6* (Fig. 2F, G). As RNAi knockdown of *RAB46* recapitulates the dysfunctional contractile vacuole phenotype of Δ*AMT1* (17), its disparate expression levels, attributable to its differential methylation, probably underpins the observed variations in the penetrance of the abnormal contractile vacuole phenotype (Fig. 2A).

### Distinct 6mA profiles in Δ*AMT6* and Δ*AMT7*

We further dissected contributions of AMT6 and AMT7 to maintenance methylation by comparing 6mA profiles in WT, Δ*AMT6*, and Δ*AMT7*. In WT, there were abundant DNA molecules containing many full-6mApT sites but few hemi-6mApT sites, while those of the opposite pattern were rare (Fig. 3A, left). By contrast, DNA in Δ*AMT7* molecules predominantly featured many hemi- 6mApT but few full-6mApT, with the opposite pattern essentially absent (Fig. 3A, right). In Δ*AMT6*, both patterns were common (Fig. 3A, middle).

**Fig. 3.**
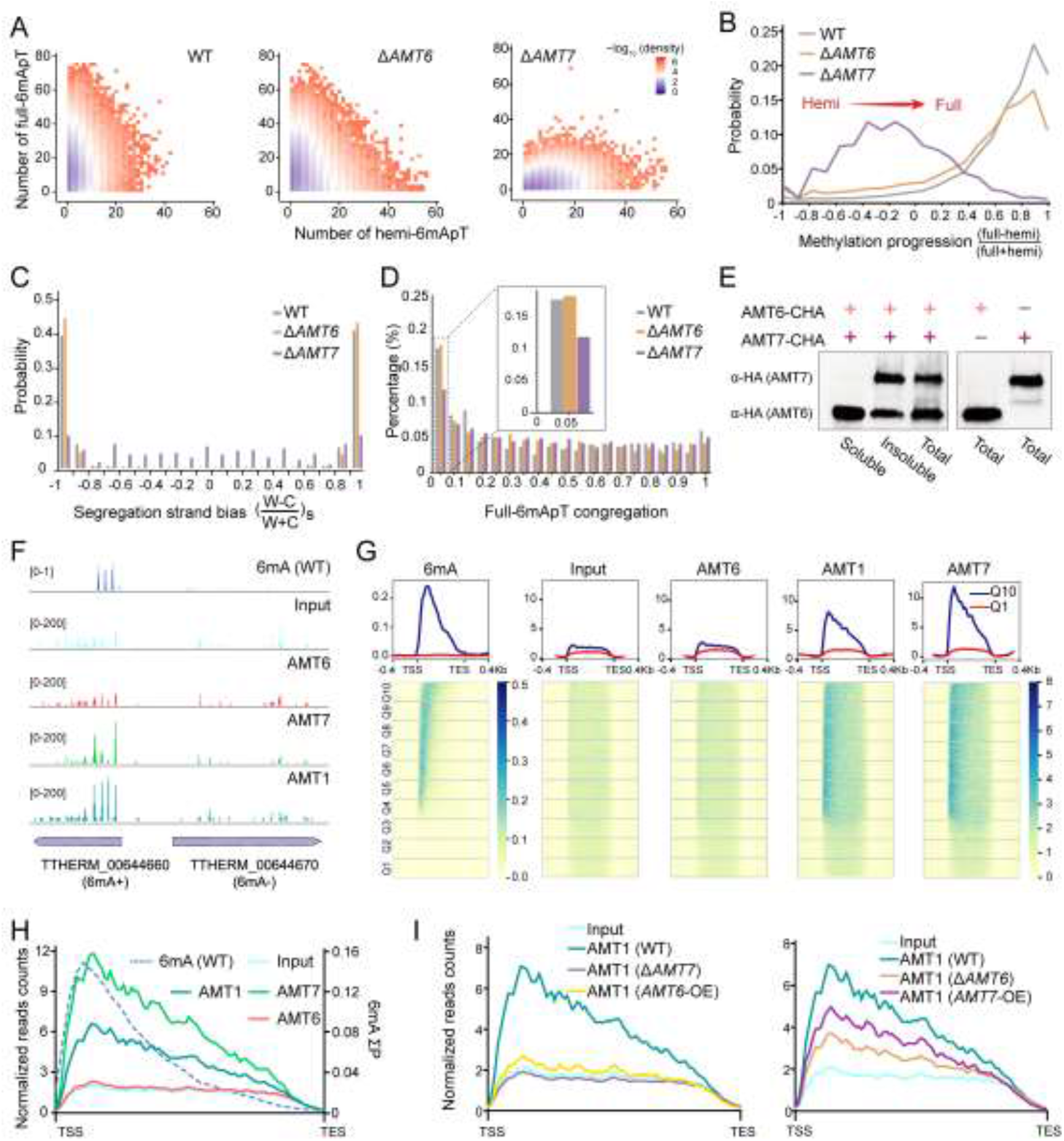
Distinct methylation characteristics, chromatin affinity, and genomic distribution of AMT6 and AMT7. A. Two-dimensional (2D) distribution of all methylated DNA molecules in WT, Δ*AMT6* and Δ*AMT7*, according to the number of hemi-6mApT (x- axis) and full-6mApT sites (y-axis) contained in each DNA molecule. B. Methylation progression (MP) in WT, Δ*AMT6* and Δ*AMT7*. MP was calculated as the difference-sum ratio between full-6mApT and hemi-6mApT on individual DNA molecules: 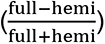. C. Segregation strand bias in WT, Δ*AMT6* and Δ*AMT7*, defined as the difference-sum ratio between hemi-6mApT sites on W and C: 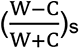. D. Full-6mApT congregation in DNA molecules undergoing hemi-to-full conversion in WT, Δ*AMT6* and Δ*AMT7*. x- axis: the probability for simulated max inter-full distances to be no greater than the observed value; y-axis: the percentage of total DNA molecules with the corresponding probability. E. Differential chromatin affinity of AMT6 and AMT7. AMT6 and AMT7 were fractionated by salt and detergent extraction (soluble and insoluble) and visualized by Immunoblot. F. A representative genomic region showing distributions of AMT1, AMT6, and AMT7 along Pol II-transcribed genes in WT. Tracks (top to bottom): 6mA penetrance, ChIP input, AMT6, AMT7, and AMT1 ChIP, gene models: 6mA+ vs. 6mA- genes. G. Distributions of 6mA, ChIP input, AMT1, AMT6, and AMT7 ChIP along the gene body in WT. The gene body, from TSS to TES, was normalized to unit length and extended in both directions by 0.4 kb. Pol II-transcribed genes (26,359 in total) were divided into 10 quantiles (Q1-Q10: low-6mA level to high-6mA level) according to their mean methylation levels in WT. AMT1, AMT6, and AMT7 distributions were normalized by input read counts of Q1 in the gene body. Composite plot showed the distribution of Q1 and Q10. Heat map displayed all 10 quantiles. H. 6mA, AMT1, AMT6, and AMT7 distributions from TSS and TES in Q10 (top quantile ranked by WT 6mA ΣP). Input was included as control for background. Y-axis: normalized read count (left scale) and 6mA ΣP (right scale). I. AMT1 distribution along the gene body was affected by changing AMT6 or AMT7 levels, showcased for Q10. Left: AMT1 distribution in WT, *AMT6*-OE, and Δ*AMT7*. Right: AMT1 distribution in WT, *AMT7*-OE, and Δ*AMT6*.

These DNA molecules were from asynchronously growing cells, representing random sampling of cell cycle progression as well as methylation progression (MP). We characterized the MP profile by focusing on DNA molecules with multiple 6mApT sites (full and hemi). In WT, most DNA molecules were at or near complete hemi-to-full conversion and dominated by full-6mApT (late MP), while few were at initial conversion stages and dominated by hemi-6mApT (early MP) (Fig. 3B). This MP profile supports that maintenance methylation is initiated shortly after DNA replication and quickly cascades to completion in WT. By contrast, there were almost no DNA molecules at late MP in Δ*AMT7* (Fig. 3B), which was not surprising given the abundance of hemi- 6mApT sites and their random distribution (Fig. 2E, Table S1). Instead, there were many DNA molecules at early MP (Fig. 3B), supporting slow methylation progression. MP distribution in Δ*AMT6* was more like WT, but less skewed towards completion (Fig. 3B). Meanwhile, there was accumulation at early MP in Δ*AMT6*, indicating a delay in initial methylation (Fig. 3B, Fig. S3A). The altered 6mA profiles in Δ*AMT6* and Δ*AMT7* reveal distinct methylation characteristics of AMT6 and AMT7 complexes.

Semiconservative 6mA transmission entails that hemi-6mApT is only found on the parental strand immediately after replication, leading to strong segregation strand bias. Indeed, for DNA molecules at early MP, hemi-6mApT sites exhibited strong segregation strand bias in Δ*AMT6* as well as in WT (Fig. 3C, Fig. S3B). However, their counterparts in Δ*AMT7* did not (Fig. 3C). This anomaly can be attributed to AMT1-independent *de novo* methylation (28), a much slower reaction that is superseded by the fast maintenance methylation in WT, and only reveals itself when maintenance methylation is dramatically reduced and delayed in Δ*AMT7*.

Many maintenance MTases (*e.g.*, *E. coli* Dam DNA MTase) are processive rather than distributive (29). We have previously showed that AMT1-dependent maintenance methylation is also processive (19). This was revealed by the congregation of full-6mApT sites in WT: for individual DNA molecules at early MP, the observed maximum distance between adjacent full-6mApT sites was often much smaller than expected (Fig. 3D, Fig. S3C). Compared to WT, full- 6mApT congregation was much reduced in Δ*AMT7*, but not in Δ*AMT6* (Fig. 3D). This supports that AMT7 complex contributes strongly to the processivity of maintenance methylation, while AMT6 complex is much less processive. This conclusion is consistent with the high chromatin affinity of the former and the low chromatin affinity of the latter (see **Distinct chromatin affinity and genomic distribution of AMT6 and AMT7**).

The percentage of total 6mApT (full + hemi, relative to all ApT) was similar in Δ*AMT7* (0.53%) and Δ*AMT1* (0.50%), even though the full-6mApT percentage was much higher in Δ*AMT7* (54.28%) than in Δ*AMT1* (3.03%) (Fig. 2B, Table S1). 6mApT levels were only slightly and uniformly increased across Pol II-transcribed genes in Δ*AMT7* relative to Δ*AMT1* (Fig. S3D). Moreover, there was a strong overlap between genomic positions methylated in Δ*AMT7* and Δ*AMT1* (Fig. S3E); in other words, the same set of ApT positions were targeted by AMT6 complex-dependent maintenance methylation (in Δ*AMT7*) and AMT1-independent *de novo* methylation (in Δ*AMT1*). These results support that in the absence of AMT7 complex, AMT6 complex-dependent maintenance methylation is incomplete, largely limited to converting hemi-6mApT deposited by AMT1-independent *de novo* methylation to full-6mApT. When comparing Δ*AMT6* or WT to Δ*AMT1*, much more substantial increases in 6mApT levels (Fig. S3D) and much less overlap between methylated genomic positions were observed (Fig. S3E). These results suggest that AMT7 complex plays a major role in maintenance methylation and targets more genomic positions—likely with contributions from its intrinsic *de novo* methylation activity.

### Distinct chromatin affinity and genomic distribution of AMT6 and AMT7

We next characterized chromatin association of AMT6 and AMT7. Under the same extraction condition, virtually all AMT7 remained chromatin-bound, while most AMT6 could be solubilized (Fig. 3E); AMT1 had a small soluble fraction and a large chromatin-bound fraction (Fig. S3K). These observations can be attributed to AMT7 complex’s higher DNA binding affinity (see **Differential activities of reconstituted AMT6 and AMT7 complexes**) and its potentially higher affinities for other chromatin associated proteins (see **AMT7 complex is recruited by transcription-associated epigenetic pathways**).

We performed chromatin immunoprecipitation followed by deep sequencing (ChIP-seq) to track genomic distribution of AMT1, AMT6, and AMT7 (Fig. 3F- H). In WT, we found that reads recovered from AMT6 ChIP were rather evenly dispersed across the gene body and essentially the same as the input (Fig. 3F-H, Fig. S3F, G), supporting that AMT6 binds chromatin with low affinity. In contrast, AMT7 ChIP reads were highly enriched towards the 5’ end of many Pol II-transcribed genes (Fig. 3F-H, Fig. S3G), a pattern reminiscent of 6mA distribution (Fig. 2C, D). This supports that AMT7 binds chromatin selectively and with high affinity. AMT1 distribution was in an intermediate state (Fig. 3F, G), closely fitted by linear combination of AMT6 and AMT7 distributions (Fig. 3H, Fig. S3G).

We then examined AMT1, AMT6, and AMT7 distribution in 10 quantiles of Pol II-transcribed genes (Fig. 3G, Fig. S3F), ranked lowest to highest by 6mA levels in WT (Fig. 2C, E). AMT7 was enriched in high-6mA genes and almost completely depleted in low-6mA genes, while AMT6 was more evenly distributed across all genes (Fig. 3G, Fig. S3F). Again, we found AMT1 distribution in an intermediate state, closely fitted by linear combination of AMT6 and AMT7 distributions (Fig. 3H, Fig. S3G, H). These results strongly suggest that AMT1 is partitioned between AMT6 complex with low chromatin affinity and AMT7 complex with high chromatin affinity.

We next examined how AMT1 distribution may be affected by changes in AMT6 or AMT7 levels (Fig. 3I, J). In addition to Δ*AMT6* and Δ*AMT7*, we also generated *Tetrahymena* strains overexpressing either AMT6 (*AMT6*-OE) or AMT7 (*AMT7*-OE) (Fig. S3I, J). In Δ*AMT7*, AMT1 was mostly found in the soluble fraction (Fig. S3K). Moreover, AMT1 distribution along the gene body was no longer highly enriched at 5’ of the gene body, even for genes with the highest 6mA levels in WT (Fig. 3I: left); instead, AMT1 was distributed in a pattern very similar to AMT6 (Fig. 3H). As mentioned above, global 6mA level was greatly reduced in Δ*AMT7* (Fig. 2E, Table S1), which may have been exacerbated by its substantial reduction in both AMT1 and AMT6 levels (Fig. S3L). Similar phenotypes were also observed upon overexpression of AMT6, including increased AMT1 solubility, fully dispersed genomic distribution, and reduced global 6mA level (Fig. 3I: left, S3K, Table S2). Note that global AMT1 level was not substantially affected in *AMT6*-OE (Fig. S3L), which may account for the less dramatic 6mA reduction than in Δ*AMT7*. These results support the repartitioning of AMT1 from AMT7 complex to AMT6 complex, which is driven by either lack of AMT7 or excess of AMT6, results in reduced chromatin affinity and specific binding of AMT1, as well as 6mA deposition.

The results were more complex when we tried to tilt the balance in the other direction. Global AMT1 level was not substantially affected upon overexpression of AMT7 (Fig. S3L). However, a substantial portion of overexpressed AMT7 was soluble (Fig. S3M) and AMT1 was also shifted from the chromatin-bound fraction to the soluble fraction (Fig. S3K), suggesting that at least some AMT7 complex was no longer tightly chromatin-bound.

Consistent with this interpretation, the ChIP result showed that AMT1 enrichment toward 5’ of the gene body was reduced (Fig. 3I: right). In Δ*AMT6*, AMT1 level remained stable, while AMT7 was substantially increased (possibly to compensate the loss of AMT6), incidentally achieving its overexpression (Fig. S3L). There was also a surprisingly large pool of AMT7 (as well as AMT1) in the soluble fraction (Fig. S3M), suggesting the presence of free AMT7 complex. Moreover, AMT1 distribution across the gene body was dispersed, but not to the extreme degree observed in Δ*AMT7* and *AMT6*-OE (Fig. 3I). Global 6mA level was not significantly affected in *AMT7*-OE and only slightly reduced in Δ*AMT6* (Fig. 2E, Table S1, S2). We attribute these observations to (near) saturation of high-affinity binding sites by AMT7 complex in WT; excess AMT7 complex was driven into the soluble fraction. This may also explain their relatively minor effects on the global 6mA level.

### AMT7 complex is connected to transcription-associated epigenetic marks

Further analysis showed that AMT7 distribution was correlated with not only 6mA, but also other transcription-associated epigenetic marks, especially the histone post-translational modification H3K4me3 and the histone variant H2A.Z (Fig. 4A-C). Pairwise comparison showed that the levels of AMT7, as well as 6mA, H3K4me3, and H2A.Z, on Pol II-transcribed genes were strongly correlated with each other (Fig. 4C). Indeed, most *Tetrahymena* genes were either enriched (triple+: 59.4%) or depleted for all three epigenetic marks (triple-: 29.0%) (Fig. S4A). Moreover, all three marks showed strong correlations with gene expression levels (Fig. 4C). As conserved transcription-associated epigenetic marks, H3K4me3 and H2A.Z are known to play important roles in transcription regulation (30–33). Our results support the same designation and functional implication for 6mA.

**Fig. 4.**
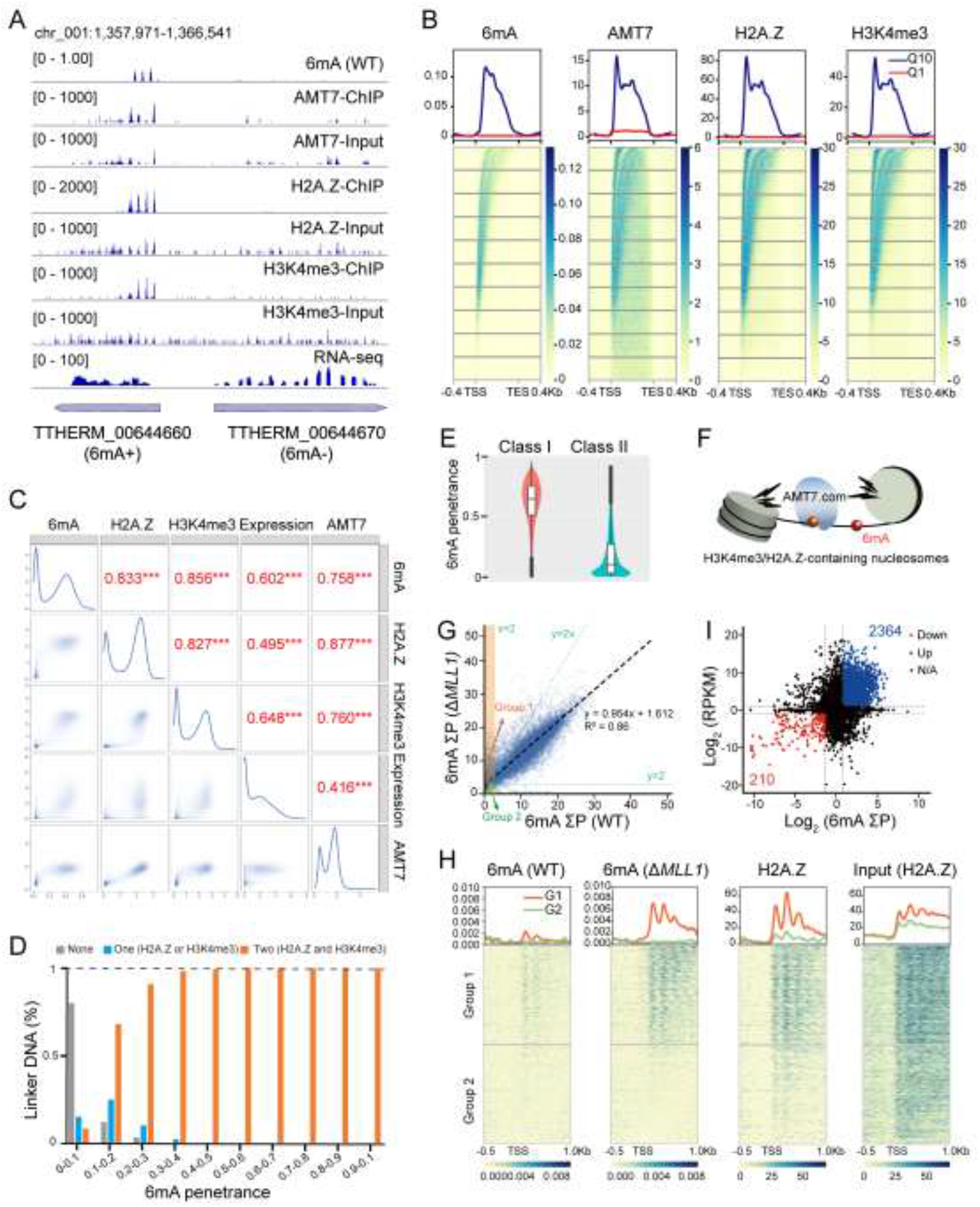
The association of AMT7 and 6mA with transcription. A. A representative genomic locus showing distributions of 6mA, AMT7, H2A.Z, and H3K4me3 as well as gene expression in WT. Note the difference in the abundance of epigenetic marks between two neighboring genes. 6mA+: high 6mA level; 6mA-: low 6mA level. B. Distributions of 6mA, AMT7, H2A.Z, and H3K4me3 along the gene body in WT. Composite plot showed the distribution of Q1 and Q10. Heat map displayed all 10 quantiles. C. Correlations between 6mA, H2A.Z, H3K4me3, AMT7, and gene expression levels. 6mA levels: 6mA ΣP. H2A.Z, H3K4me3, and AMT7 levels: ChIP reads normalized by input across the gene body. Gene expression levels: RPKM for RNA-seq of WT. D. 6mA penetrance in linker DNA (LD) dependent on whether none, one, or two of the flanking nucleosomes were enriched for H2A.Z and/or H3K4me3. 6mA was divided into 10 groups according to their penetrance (x-axis). The percentage of a specific type of LD in each 6mA group was calculated (y-axis). E. Higher 6mA penetrance in LD when each of the two flanking nucleosomes were enriched for both H2A.Z and H3K4me3 (Class I) than when one mark was missing from either nucleosome or both (Class II). F. Model: AMT7 complex is targeted by the dinucleosome containing H2A.Z and H3K4me3 for efficient 6mA deposition on LD. G. 6mA levels of individual genes in WT and Δ*MLL1*. In Δ*MLL1*, 6mA levels substantially increased in many WT low-6mA genes, while remained stable in most WT high-6mA genes (dotted diagonal line). Each gene was assigned a coordinate: 6mA ΣP for WT (x-axis) and Δ*MLL1* (y-axis). Group 1 were genes with low 6mA level in WT but elevated 6mA level in Δ*MLL1*: ΣP(Δ*MLL1*)≥ 2*ΣP(WT), ΣP(Δ*MLL1*)≥ 2, ΣP(WT)≤ 2. Group 2 were genes with stably low 6mA level in both cells: ΣP(Δ*MLL1*)< 2*ΣP(WT), ΣP(Δ*MLL1*)< 2, ΣP(WT)≤ 2. H. 6mA in Δ*MLL1* spread to genomic regions moderately enriched for H2A.Z (G1, group 1 defined in G), but remained unaffected in regions devoid of H2A.Z (G2, group 2 defined in G). I. Relationship between changes in 6mA and gene expression in WT and Δ*MLL1*. Δ*MLL1* versus WT: Log2(FoldChange) for individual genes in 6mA (x-axis: ΣP) and expression (y-axis: RPKM). Blue: genes with significantly increased 6mA and expression levels (>2×). Red: genes with significantly decreased 6mA and expression levels (<0.5×).

AMT7, 6mA, H3K4me3, and H2A.Z were all co-enriched towards the 5’ end of Pol II-transcribed genes (Fig. 4A, B), consistent with our previous reports (20). AMT7 was enriched on nucleosomes with high levels of H3K4me3 or H2A.Z (Fig. S4B). 6mA levels were the highest for linker DNA flanked on both sides by such nucleosomes, much reduced if only flanked on one side, and minimal if not flanked at all (Fig. 4D). Among linker DNA flanked by two such nucleosomes, 6mA levels were high if both H3K4me3 and H2A.Z were enriched in both nucleosomes, but much reduced if either mark was depleted from either nucleosome (Fig. 4E). In addition, 6mA penetrance dropped precipitously in linker DNA significantly exceeding the regular length (Fig. S4C, D). All these observations suggest that ***1)*** H3K4me3/H2A.Z-containing nucleosomes represent high-affinity sites for AMT7 complex; and ***2)*** a pair of such nucleosomes, joined by 6mA-decorated linker DNA of proper length, is a basic unit for AMT7 complex-dependent 6mA transmission (Fig. 4F).

After establishing correlations between AMT7 and the triple marks in WT, we next investigated their response to perturbations. As H2A.Z is encoded by an essential gene in *Tetrahymena* (34), we focused on loss-of-function analyses of H3K4me3. In Δ*MLL1*, knocking out the only *Tetrahymena* homologue of mixed lineage leukemia (MLL/KMT2) family histone MTases (35), H3K4me3 global levels were greatly diminished (Fig. S4E, F). While 6mA global level and full-6mApT percentage in Δ*MLL1* were comparable to WT (Fig. S4G, H, Table S1), there were notable alterations to 6mA patterns (Fig. 4G, Fig. S3A- C). DNA molecules at early MP accumulated (Fig. S3A), but the delay of initial methylation was not severe enough to affect the segregation strand bias of hemi-6mApT (Fig. S3B).

6mA distribution across Pol II-transcribed genes was also altered in Δ*MLL1*. For high-6mA genes in WT, their 6mA levels were mostly unaffected; however, for low-6mA genes in WT, their 6mA levels were often substantially increased (Fig. 4G, H, Fig. S4I-K). Furthermore, while full-6mApT percentage was slightly reduced in Δ*MLL1* relative to WT for 6mA+ genes (78.01% and 80.74%, respectively), it was increased for 6mA- genes (58.55% and 41.38%, respectively) (Table S3). This was not caused by random increases of methylation background in Δ*MLL1*: in Pol I-transcribed rDNA, 6mA remained at the minimum level (Table S3); in genes with induced 6mA, 6mA distribution also oscillated with the 200-bp periodicity and exhibited a strong bias toward 5’ end of the gene body (Fig. 4H, Fig. S4L). Therefore, while H3K4me3 is not required for 6mA (possibly due to redundancy in AMT7 complex recruitment mechanisms), it plays a crucial role in limiting off-target 6mA, likely via its contribution in recruiting and sequestering AMT7 complex. Intriguingly, genes with induced 6mA in Δ*MLL1* often had higher H2A.Z levels than those that remained 6mA-low (Fig. 4H), implicating that even the off-target 6mA sites were not random, but rather biased towards genes decorated with H2A.Z.

We also examined gene expression changes in response to perturbations. Despite deficiency in the euchromatic mark H3K4me3, many more genes were significantly up-regulated than down-regulated in Δ*MLL1* relative to WT (Fig. 4I, Fig. S4M). This was especially prominent for 6mA- genes, many of which were expressed at the minimum level in WT but were induced in Δ*MLL1*, accompanied by significantly increased 6mA levels; their changes in gene expression levels showed positive correlations with changes in 6mA levels (Fig. 4I). In contrast, for many 6mA+ genes in WT, they showed minimum 6mA in Δ*AMT7*, while their expression was mostly reduced; genes with less extreme reduction in 6mA levels showed more divergent response in expression levels (Fig. S4N). These results suggest that for many Pol II-transcribed genes, a threshold 6mA level may be required for their expression.

## Discussion

### AMT6 and AMT7 complexes achieve specific methylation via multiple recognitions

In *Tetrahymena*, 6mA is transmitted in an AMT1-dependent semiconservative manner (19). Here we characterize two AMT1 complexes, with AMT6 and AMT7 as their mutually exclusive components. Both AMT6 and AMT7 complexes prefer hemi-methylated over unmethylated ApT sites as the substrate, consistent with their roles in maintenance methylation. However, they are recruited by different protein-protein interactions (PPI) and feature distinct MTase activities.

AMT7 complex targeting is mediated by transcription-associated epigenetic pathways. Here we highlight H2A.Z as a nexus of these interactions. H2A.Z is enriched downstream of transcription start sites (TSS), where it is associated with active epigenetic marks like H3K4me3 and histone acetylation (21, 36–41). In *Tetrahymena*, most genes are either simultaneously marked by 6mA, H2A.Z, and H3K4me3 (triple+: 59.4%), or do not have any (triple-: 29.0%). Therefore, the triple marks may underpin the transcription-associated epigenetic landscape. We propose that a pair of H3K4me3/H2A.Z-containing nucleosomes, joined by 6mA-decorated linker DNA of proper length, represent a high-affinity site for AMT7 complex and a basic unit for AMT7 complex-dependent 6mA transmission. It is worth noting that AMT7 complex remains sequestered at these high-affinity sites even after completion of maintenance methylation. This may help to prevent non-specific methylation across the rest of the genome.

### AMT6 and AMT7 complexes coordinate in maintenance methylation

Our findings support the existence of two AMT1 complexes that are disparate in their chromatin affinities and MTase activities. AMT7 complex is stably associated with chromatin, processive, and responsible for the bulk of 6mA deposition. In contrast, AMT6 complex is more soluble, less processive, and important for expediting maintenance methylation, especially at the initial stage. We attribute the high affinity of AMT7 complex to its strong binding of H3K4me3/H2A.Z-containing nucleosome. AMT7 complex may directly engage the histones, or more likely, be indirectly recruited by the protein-protein interactions we have identified. AMT7 complex also tightly binds dsDNA, which may involve a stable clamp-like structure wrapped around linker DNA. Here we distinguish two separate steps for specific targeting of AMT6 and AMT7 complexes. The initial step recruits AMT6 or AMT7 by PPI, while the subsequential step leads to stable binding and methylation of linker DNA. Their apparent difference in chromatin affinity results from differential kinetics in these two steps.

AMT6 complex and AMT7 complex compete for the same pool of AMT1 (most likely in the form of AMT1 subcomplex); in other words, AMT1 actively exchanges between these two complexes. Our perturbation analyses support that AMT6 and AMT7 levels in WT *Tetrahymena* are (near) optimal for 6mA maintenance methylation. AMT7 is expressed at a level to (nearly) saturate high-affinity sites, but not in excess to drive AMT7 complex into the soluble fraction. AMT6 is expressed at a level to expedite maintenance methylation, but not in excess to compete away AMT7 complex binding at high-affinity sites. Maintenance methylation is negatively affected by deletion or overexpression of both AMT6 and AMT7. This finetuning also suggests that the balance between the two MTase complexes is tightly regulated and functionally relevant.

AMT6 complex alone is limited to converting hemi-6mApT deposited by *de novo* methylation to full-6mApT. AMT7 complex alone can substantially expand 6mA repertoire and increase 6mA levels on Pol II-transcribed genes, but is delayed in methylation progression. These results strongly suggest that in WT *Tetrahymena*, AMT6 and AMT7 co-occupy target sites, possibly by sharing AMT1 subcomplex. Mediated by multiple recognitions, AMT6 and AMT7 complexes coordinate in maintenance methylation, to achieve specific, efficient, and robust transmission of 6mA as a eukaryotic epigenetic mark.

## Materials and Methods

### Cell culture

*Tetrahymena thermophila* wild-type (WT) strains (SB210 and CU428) were obtained from the *Tetrahymena* Stock Center (http://tetrahymena.vet.cornell.edu). Homozygous homokaryon strain Δ*AMT1* and somatic hemagglutinin (HA) tagged strains (AMT1-NHA and AMT7-CHA) were generated in our previous studies (19, 20). Cells were grown in SPP medium at 30°C (42, 43).

### Immunoprecipitation (IP) and mass spectrometry analysis

Approximately 5 × 10^7^ purified MACs were solubilized with 500 U Benzonase (Millipore, 712053N) and incubated with pre-washed HA beads (Sigma, E6779). Targeted proteins were detected by mass spectrometry in Analysis Center of Agrobiology and Environmental Sciences, Zhejiang University.

### UHPLC-QQQ-MS/MS analysis

Genomic DNA was digested following established protocols (20). Samples were analyzed by ultra-high-performance liquid chromatography-tandem mass spectrometry (UHPLC-QQQ-MS/MS) (13, 44). The selective MRM transitions were detected under m/z 266/150 for 6mA and m/z 252/136 for dA. The ratio of 6mA/A was quantified by the calibration curves of nucleoside standards running at the same time.

### Recombinant protein purification

Full length AMT1, AMT6, AMT7, AMTP1, AMTP2 ORFs were codon-optimized for bacterial expression. *E. coli* was cultured to an optical density (OD600) of 0.8 at 37°C and protein expression was induced by 0.25 mM IPTG at 16°C overnight. Proteins were first purified by Ni-NTA (Novagen, USA) affinity chromatography and eluted through on-column tag cleavage by ULP1 protease. The eluate was further purified by passage through a Heparin ion-ex-change column (GE Healthcare, USA) and a Superdex 200 10/300 Increase column (GE Healthcare, USA).

### Methyltransferase assay

For non-radioactive MTase assay, 2 μM protein was mixed with 1μg pUC19 plasmid purified from *dam-*/*dcm- E. coli*, 10 μM SAM in 40 μL reaction buffer. Samples were incubated at 30°C for 30 min and inactivated at 65°C for 20 min. Methylated DNA samples were treated with *Dpn*I at 37°C for 30 min. For radioactive MTase assay, 0.05 μM protein was mixed with 0.5 μM [^3^H]SAM and dsDNA at various concentrations. Samples were incubated at 30°C for 30 min and subsequently spotted onto Hybond-XL membrane (GE Healthcare). Each membrane was immersed in Ultima Gold (PerkinElmer) and used for scintillation counting on a PerkinElmer MicroBeta2 (PerkinElmer).

### Electrophoretic mobility shift assay (EMSA)

EMSA was performed using dsDNA and proteins purified as above. To compare substrate preference, 4 μM protein was incubated with 4 μM different DNA substrates. Reactions were run on a 6% native polyacrylamide gel (acrylamide:bis-acrylamide=37.5:1) in 1 x TBE buffer at 100 V for 30 min. The gel was stained with ethidium bromide.

### Surface plasmon resonance (SPR) measurement

Biotinylated-A/6mA DNA probes were captured on SA sensor chip (∼100 response units). SPR experiments were performed using Biacore T200 instrument (Cytiva). Proteins with increasing concentrations were injected into the A/6mA probes surface for 60 s at a flow rate of 30 μL/min and dissociated for 90 s (AMT6 complex) or 480 s (AMT7 complex) in running buffer. Binding data were collected and analyzed by Biacore™ Insight Evaluation Software.

### RNA-seq and ChIP-seq data analysis

RNA samples and ChIP samples were sequenced by Illumina HiSeq platform with 150 bp dual end sequencing. For RNA-seq, differentially expressed genes (DEGs) were identified by DESeq2 (Log2(FoldChange) > 1 or < -1, *P* < 0.05). For ChIP-seq, the MACs of HA-tagged strains were purified and digested according to established protocols (45). Well-annotated Pol II-transcribed genes (26,359 in total) were scaled to unit length and divided into 10 quantiles (Q1-Q10: low-6mA level to high-6mA level) according to their methylation levels per unit length. Each group from Q1 to Q10 was normalized using the area under the curve (AUC) from TSS to TES of the Q1 group. Enriched genes were defined as ChIP/input ratio higher than 1 (normalized by total counts respectively).

### SMRT sequencing and data analysis

SMRT sequencing data was analyzed following previously described procedures, including gene methylation level analysis, methylation progress, segregation strand bias, full-6mApT congregation, autocorrelation and cross- correlation analysis (19).

### Phylogenetic analysis

AMT6 and AMT7 of *T. thermophila* were used as seed sequences to search for respective orthologues in other eight species of the *Tetrahymena* genus in the *Tetrahymena* comparative genome database (TCGD, http://ciliate.ihb.ac.cn/tcgd/) using BLASTP (46). All candidates were aligned using mafft (47) in G-INS-I model and phylogenetic trees were constructed using IQ-TREE2 (48) under default parameters expect for LG+C20 model and 1,000 bootstraps (-m LG+C20 -B 1000).

## Data availability

The latest SB210 MAC genome can be found at the *Tetrahymena* genome database (TGD) (http://ciliate.org) (49, 50).

## Acknowledgement

This work was supported by the National Natural Science Foundation of China (32125006, 32070437 to SG, 32200399 to YW), Natural Science Foundation of Shandong Province of China (ZR2021QC046 to YW), the Fundamental Research Funds for the Central Universities (202441014 to YW), and Department of Biochemistry and Molecular Medicine at University of Southern California (YL). The authors would like to thank the following people and labs for assistance with this study: Dr. Kensuke Kataoka (National Institute for Basic Biology, Japan) for sharing the 3×G196 construct and Mr. Zhaorui Zhou (Ocean University of China, OUC) for sharing the methods of phylogenetic tree construction. Our special thanks are given to Dr. Weibo Song (OUC) for his helpful suggestions during drafting the manuscript. SMRT sequencing was performed at the Genomics Core Facility, Icahn School of Medicine at Mount Sinai and Novogene Co., Ltd. (Beijing, China). High-performance computing resources for data processing were provided by Marine Big Data Center of Institute for Advanced Ocean Study at OUC and the Center for Advanced Research Computing (CARC) at the University of Southern California.

## Author contribution

YW, YL and SG conceived the study and designed the experiments; YW performed most of the experiments; BN reconstituted AMT1 complex and studied its enzyme kinetics; FY and WY performed the bioinformatic analysis with BP; ZZ generated and characterized the related *Tetrahymena* strains with AJ; JN and WZ performed the mass spectrometry analysis; YW, YL and SG wrote the paper, with inputs from all authors. All authors read and approved the final manuscript.

The authors declare no conflict of interest.

