## Supplemental Figures and Tables for "Dual modes of DNA N^6^-methyladenine maintenance by distinct methyltransferase complexes"

#These authors contributed equally

### Supplemental figures S1-S4

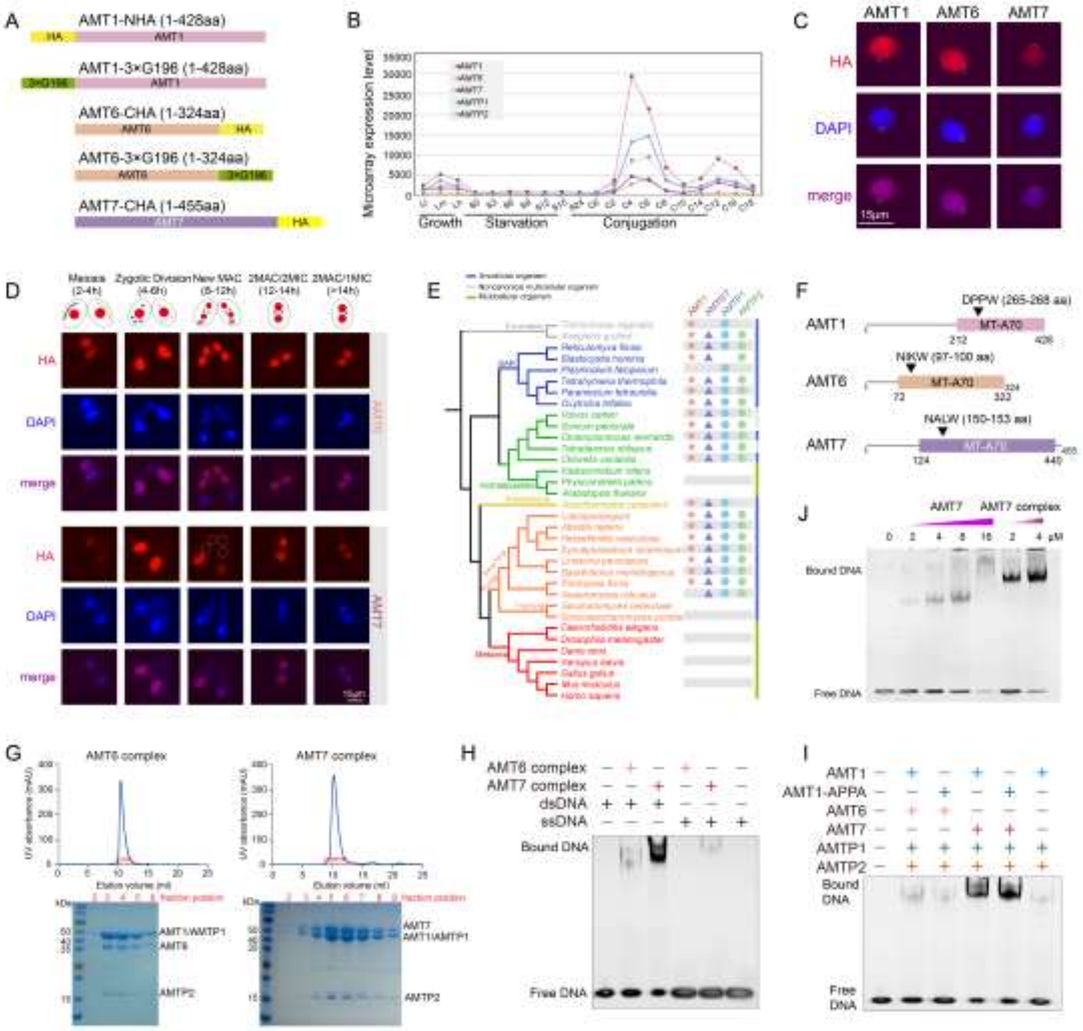


**Fig. S1. Two distinct AMT1 complexes with AMT6 or AMT7 as their mutually exclusive components.**

A. Schematics of HA or 3×G196- epitope-tagging for components of AMT1 complexes.

B. Gene expression levels of *AMT1*, *AMT6*, *AMT7*, *AMTP1*, and *AMTP2* at different physiological stages. Microarray data were downloaded from *Tetrahymena* Functional Genomics Database (1).

C. Cellular localization of AMT6 and AMT7 in growing cells. Both proteins were HA-tagged and detected by an α-HA antibody. Micronucleus (MIC) was outlined (dotted circles) without detectable HA signals.

D. Dynamic distribution of AMT6 and AMT7 during conjugation. AMT6 and AMT7 were HA-tagged for both somatic and germline copies. Note the absence of AMT7 signal during the new MAC stage (dotted circles). Conjugation stages were monitored according to nuclear events (see schematics on the top).

E. Phylogenetic distributions of conserved core components for AMT1 complexes, modified from our previous result (2).

F. Conservation of the catalytic motif in MT-A70 domains of AMT1 (DPPW), AMT6 (NIKW), and AMT7 (NALW).

G. Gel filtration chromatography (top panel) and SDS-PAGE (bottom panel) showing co-fractionation of the components of AMT6 (left) or AMT7 (right) complexes.

H. Electrophoretic Mobility Shift Assay (EMSA) showing DNA binding affinity of two complexes. Both complexes showed a preference towards double-stranded DNA (dsDNA) over single-stranded DNA (ssDNA).

I. Electrophoretic Mobility Shift Assay (EMSA) showing specific binding of dsDNA for AMT6 and AMT7 complexes, AMT1 mutation (AMT1-APPA) retaining the DNA binding affinity (lane 3 vs. lane 2; lane 5 vs. lane4), and weak binding for AMT1 subcomplex (similar to that of AMT6 complex) (lane 6 vs. lane 2).

J. EMSA showing that AMT7 alone did not bind dsDNA tightly.

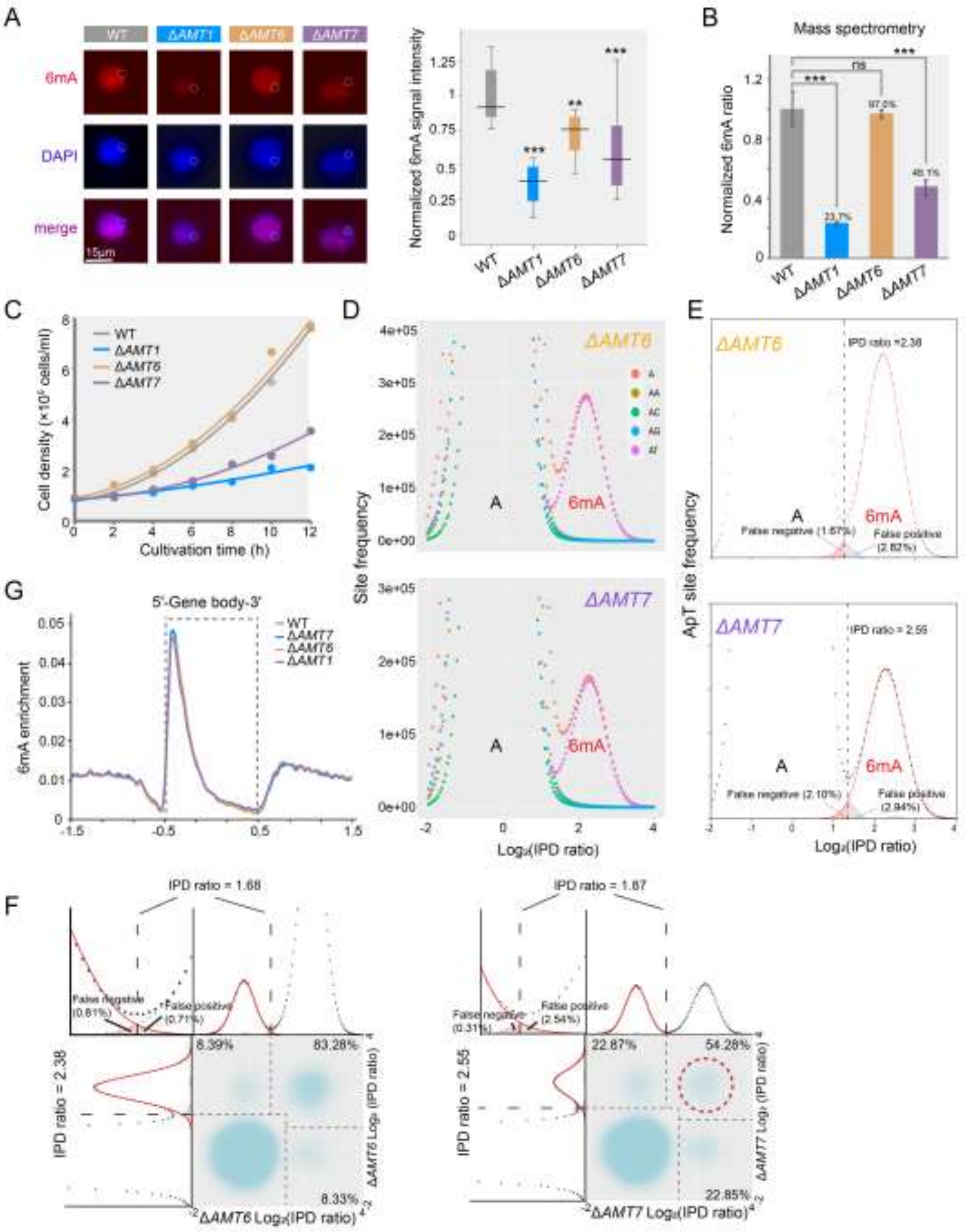


**Fig. S2. Distinct phenotypes of Δ*AMT6* and Δ*AMT7*.**

A. IF staining of 6mA and statistical analysis of IF signal intensity in vegetative WT, Δ*AMT1*, Δ*AMT6* and Δ*AMT7*. Note the absence of 6mA signal in MICs (dotted circles). Statistical analysis was performed with random images of each cell (n=261 (WT), 283 (Δ*AMT1*), 358 (Δ*AMT6*), 178 (Δ*AMT7*)). ** $P < 0.01$; *** $P < 0.001$.

B. Mass spectrometry analysis of 6mA. Data were presented as mean ± S.D. Student's *t*-test was performed. *** $P < 0.001$; ns: not significant ($P > 0.05$).

C. Growth rates of WT and knockout cells. Cells were counted using a Coulter counter at indicated time points.

D. 6mA exclusively present in the ApT dinucleotide in Δ*AMT6* and Δ*AMT7*. IPD ratio distribution ($Log_2$) of all A sites in Δ*AMT6* and Δ*AMT7* was plotted in the ApA, ApC, ApG, and ApT dinucleotide, respectively.

E. Deconvolution of the 6mA peak and the unmodified A peak for IPD ratio distributions ($Log_2$) in the ApT dinucleotide. Note the low false positive and false negative rates of 6mA calling in Δ*AMT6* and Δ*AMT7*.

F. Demarcation of the four methylation states of ApT duplexes in Δ*AMT6* and Δ*AMT7* by their IPD ratio (IPDr) on W and C, respectively. For bulk ApT duplexes, the IPDr threshold for 6mA calling was set (2.38 for Δ*AMT6* and 2.55 for Δ*AMT7*) according to deconvolution based on Gaussian fitting of the small 6mA peak. For ApT duplexes with one 6mA, the IPDr threshold for calling 6mA on the opposite strand was set (1.68 for Δ*AMT6* and 1.87 for Δ*AMT7*) according to deconvolution based on Gaussian fitting of the small unmodified A peak.

G. 6mA distribution towards the 5' end of gene body of Pol II-transcribed genes in WT, Δ*AMT1*, Δ*AMT6*, and Δ*AMT7*. Genes were scaled to the same area under the curve from TSS to TES. Y axis represented 6mA frequency (6mA amount at a certain position/total 6mA amount).

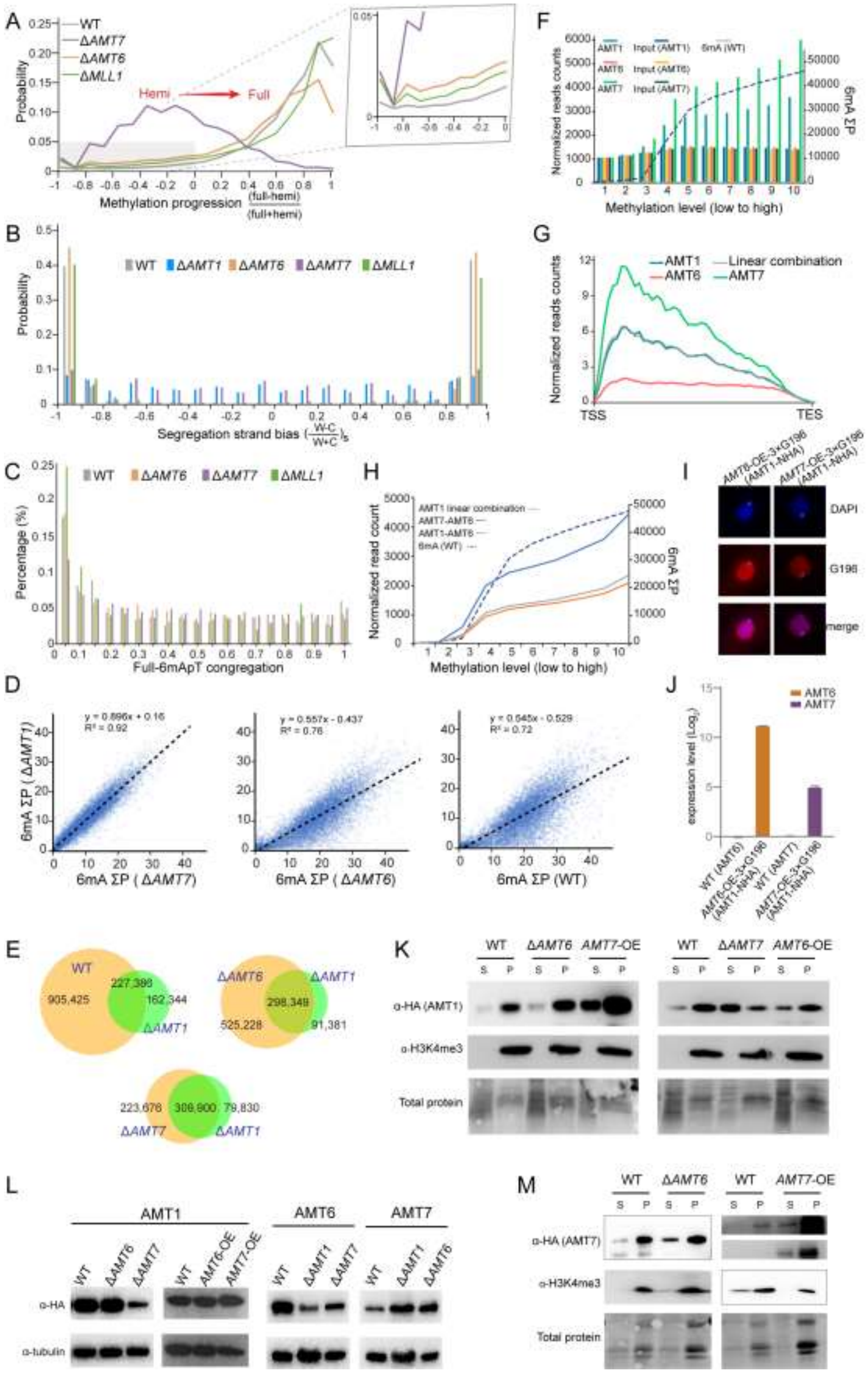

A
Hemi
Full
Methylation progression (full-hemi)/(full+hemi)
Probability
B
Segregation strand bias ((W-C)/(W+C))s
C
Full-6mApT congregation
Percentage (%)
D
y = 0.896x + 0.16
R² = 0.92
y = 0.557x - 0.437
R² = 0.76
y = 0.545x - 0.529
R² = 0.72
6mA ΣP (ΔAMT1)
6mA ΣP (ΔAMT7)
6mA ΣP (ΔAMT6)
6mA ΣP (WT)
E
905,425
227,386
162,344
525,228
298,349
91,381
223,676
309,900
79,830
F
Methylation level (low to high)
Normalized reads counts
6mA ΣP
G
AMT1
AMT6
Linear combination
AMT7
Normalized reads counts
TSS
TES
H
AMT1 linear combination
AMT7-AMT6
AMT1-AMT6
6mA (WT)
Normalized read count
Methylation level (low to high)
I
DAPI
G196
merge
J
expression level (Log2)
K
α-HA (AMT1)
α-H3K4me3
Total protein
L
AMT1
AMT6
AMT7
α-HA
α-tubulin
M
α-HA (AMT7)
α-H3K4me3
Total protein

**Fig. S3. Distinct methylation characteristics, chromatin affinity, and genomic distribution of AMT6 and AMT7.**

A. Methylation progression (MP) in WT, Δ*AMT6*, Δ*AMT7*, and Δ*MLL1*. MP was calculated by the difference-sum ratio between full-6mApT and hemi-6mApT on individual DNA molecules: $(\frac{\text{full}-\text{hemi}}{\text{full}+\text{hemi}})$. Note that DNA molecules were accumulated in early methylation progress in Δ*AMT6*, Δ*AMT7* and Δ*MLL1*, as shown in the zoom-in plot.

B. Segregation strand bias in WT, Δ*AMT1*, Δ*AMT6*, Δ*AMT7*, and Δ*MLL1*, defined as the difference-sum ratio between BrdU sites on W and C: $(\frac{\text{W}-\text{C}}{\text{W}+\text{C}})_S$.

C. Full-6mApT congregation in DNA molecules undergoing hemi-to-full conversion in WT, Δ*AMT6*, Δ*AMT7*, and Δ*MLL1*. The maximum observed distance between adjacent full-6mApT sites (max inter-full distances) was much smaller than expected, and as a result rarely represented (probability ≤ 0.05) in simulated controls, in which full-6mApT and hemi-6mApT positions were randomly permutated. In other words, there was a strong tendency for multiple maintenance methylation events to occur in nearby ApT dinucleotides. X- axis: the probability for simulated max inter-full distances to be no greater than the observed value; y-axis: the percentage of total DNA molecules with the corresponding probability.

D. 6mA levels of individual genes in WT, Δ*AMT1*, Δ*AMT6* and Δ*AMT7*. Each gene was assigned a coordinate: sum of 6mA penetrance values for all methylated ApT positions in the gene body (ΣP) for WT, Δ*AMT6* and Δ*AMT7* (x-axis) vs. Δ*AMT1* (y-axis).

E. Overlap between methylated ApT positions in WT, Δ*AMT6* and Δ*AMT7* compared individually with Δ*AMT1*. 6mA ApT number ≥3 for any ApT positions in each sample.

F. AMT1, AMT6, and AMT7 distributions in 10 quantiles of genes ranked from low to high according to their 6mA levels (ΣP) in WT. Input was included as control for background. Y-axis: normalized read count (left) and 6mA ΣP (right).

G. AMT1 distribution as the linear combination of AMT6 and AMT7 distributions along the gene body, showcased in Q10 (top quantile ranked by WT 6mA levels). The linear combination of AMT6 and AMT7 minimized the sum of squared errors when fitting the result to AMT1 distribution.

H. AMT1 distribution as the linear combination of AMT6 and AMT7 distributions in 10 quantiles of genes ranked from low to high according to their 6mA levels (ΣP) in WT. The ChIP-reads of AMT1, AMT7 and AMT1 liner combination was all normalized by AMT6 ChIP-reads. Note that the distribution curve of AMT7 fitted well with that of 6mA. Y-axis: normalized read count (left) and 6mA ΣP (right).

I. IF staining showing that ectopically expressed AMT6 and AMT7 were specifically located in the MAC, but absent in the MIC (dotted circles). Cells were incubated with 1.5 μg/mL cadmium chloride for 4-5 h before fixed for staining.

J. Expression level of *AMT6* and *AMT7* in *AMT6*-OE-3×G196 (AMT1-NHA) and *AMT7*-OE-3×G196 (AMT1-NHA) were largely upregulated, compared to that in WT.

K. Differential chromatin affinity of AMT1 in WT, *AMT6*-OE, *AMT7*-OE, Δ*AMT6*, and Δ*AMT7*. H3K4me3 was used as the loading control of chromatin-bound fraction. Total protein was detected by No-Stain™ Protein Labeling Reagent (Invitrogen, A44449). Note the soluble fraction (S) and the chromatin-bound fraction (P).

L. Immunoblot showing global levels of AMT1, AMT6, and AMT7, in WT, *AMT6*-OE, *AMT7*-OE, Δ*AMT6*, and Δ*AMT7*. Tubulin was used as the internal control.

M. Differential chromatin affinity of AMT7 in WT, *AMT7*-OE, and Δ*AMT6*. H3K4me3 was used as the loading control of chromatin-bound fraction. Total protein was detected by No-Stain™ Protein Labeling Reagent (Invitrogen, A44449). Note the soluble fraction (S) and the chromatin-bound fraction (P).

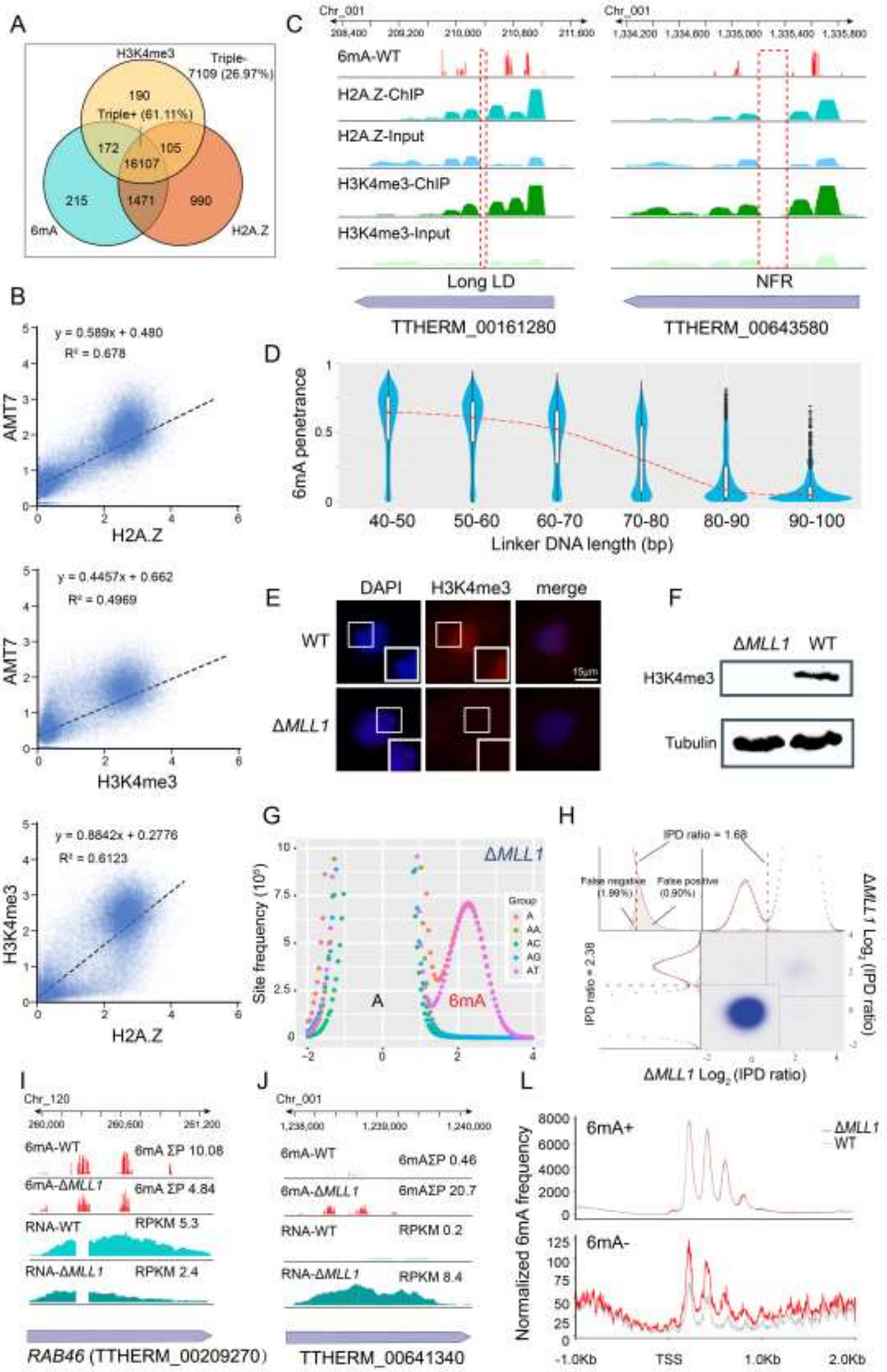
A
H3K4me3
Triple-
7109 (26.97%)
190
Triple+ (61.11%)
172
105
16107
215
1471
990
6mA
H2A.Z
B
y = 0.589x + 0.480
R² = 0.678
AMT7
H2A.Z
y = 0.4457x + 0.662
R² = 0.4969
AMT7
H3K4me3
y = 0.8842x + 0.2776
R² = 0.6123
H3K4me3
H2A.Z
C
Chr_001
6mA-WT
H2A.Z-ChIP
H2A.Z-Input
H3K4me3-ChIP
H3K4me3-Input
Long LD
NFR
TTHERM_00161280
TTHERM_00643580
D
6mA penetrance
40-50
50-60
60-70
70-80
80-90
90-100
Linker DNA length (bp)
E
DAPI
H3K4me3
merge
WT
ΔMLL1
15μm
F
ΔMLL1
WT
H3K4me3
Tubulin
G
Site frequency (10⁵)
ΔMLL1
Group
A
AA
AC
AG
AT
A
6mA
H
IPD ratio = 1.68
False negative (1.99%)
False positive (0.90%)
IPD ratio = 2.38
ΔMLL1 Log₂ (IPD ratio)
ΔMLL1 Log₂ (IPD ratio)
I
Chr_120
6mA-WT
6mA ΣP 10.08
6mA-ΔMLL1
6mA ΣP 4.84
RNA-WT
RPKM 5.3
RNA-ΔMLL1
RPKM 2.4
RAB46 (TTHERM_00209270)
J
Chr_001
6mA-WT
6mAΣP 0.46
6mA-ΔMLL1
6mAΣP 20.7
RNA-WT
RPKM 0.2
RNA-ΔMLL1
RPKM 8.4
TTHERM_00641340
L
Normalized 6mA frequency
6mA+
ΔMLL1
WT
6mA-
-1.0Kb
TSS
1.0Kb
2.0Kb

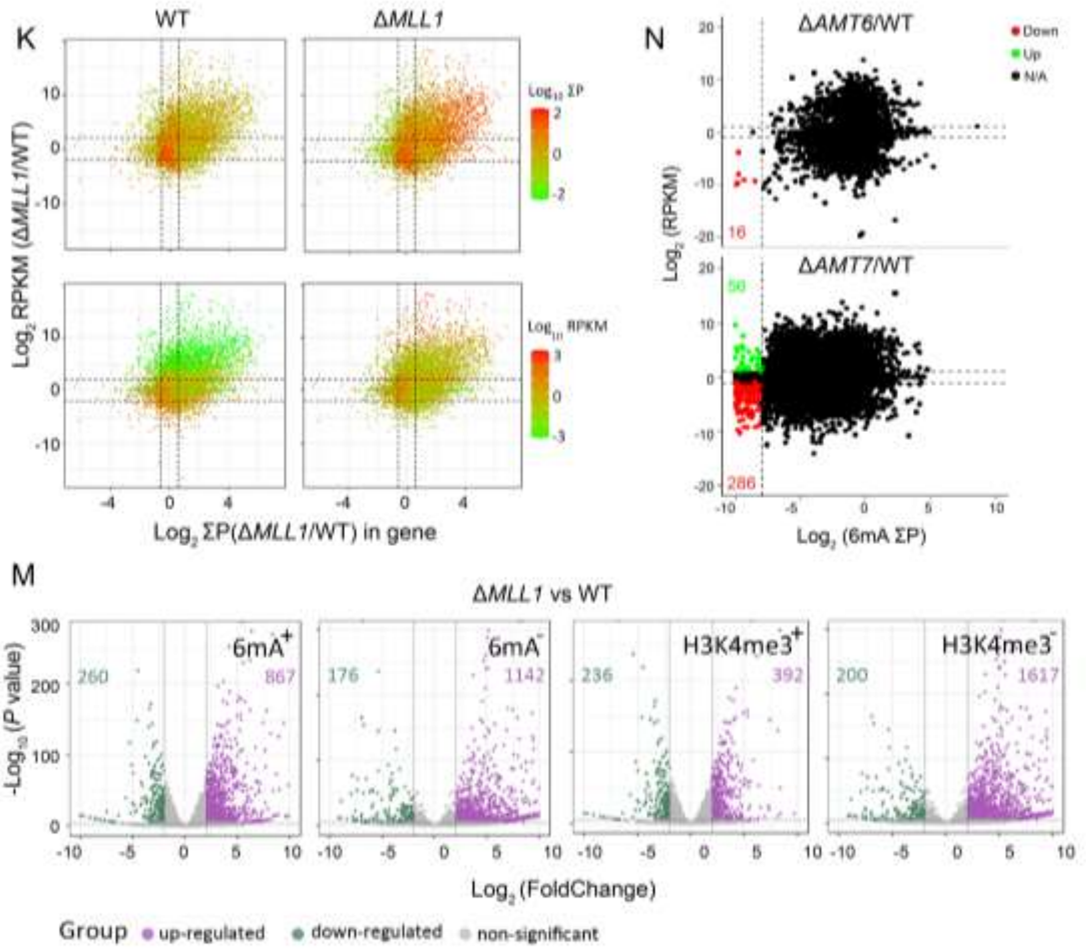


**Fig. S4. The association of AMT7 and 6mA with transcription.**

A. Strong overlap between 6mA+, H2A.Z+, and H3K4me3+ genes (triple+). Genes enriched for all or none of three marks: 6mA, H2A.Z, H3K4me3 were designated as triple+ or none triple-, respectively. Numbers and percentages (paratheses) of well-annotated *Tetrahymena* genes in each category were shown.

B. Strong pairwise correlations of nucleosomes associated with AMT7, H2A.Z and H3K4me3. For levels of AMT7, H2A.Z, and H3K4me3, ChIP/input values were calculated over the gene body.

C. 6mA depletion in a long linker DNA (LD) and a nucleosome-free region (NFR), even if their flanking nucleosomes were enriched for H3K4me3 and H2A.Z.

D. 6mA levels (mean penetrance) in linker DNA dependent on its length.

E. H3K4me3 abolished in Δ*MLL1*, as shown by immunofluorescence (IF) staining.

F. H3K4me3 abolished in Δ*MLL1*, as shown by immunoblotting. α-tubulin was used as the loading control.

G. IPD ratio distribution ($Log_2$) of adenine sites in Δ*MLL1* (left). Deconvolution of the 6mA peak and the unmodified A peak for IPD ratio distributions ($Log_2$) in the ApT dinucleotide (right).

H. Demarcation of the four methylation states of ApT duplexes by their IPD ratios on W and C in Δ*MLL1*.

I. High 6mA and expression levels of *RAB46* in WT and Δ*MLL1*.

J. A representative gene with induced 6mA and expression levels in Δ*MLL1* relative to WT.

K. Relationship between changes in 6mA and gene expression. ΔMLL1 versus WT: $Log_2$(FoldChange) for individual genes in 6mA (x: ΣP) and expression (y: RPKM). Color scales represent individual genes' 6mA (top) and expression levels (bottom) in WT (left) and Δ*MLL1* (right).

5.

A. 6mA distribution in 6mA+ and 6mA- genes of WT and Δ*MLL1*. Pol II-transcribed genes were aligned at their transcription start sites (TSS) and extended upstream (-1000 bp) and downstream (2000 bp). X axis: distance from TSS. Y axis: cumulative 6mA penetrance (WT: gray; Δ*MLL1*: red). 6mA+ (top) and 6mA- genes (bottom) were plotted separately. Note that for 6mA+ genes, the WT and Δ*MLL1* curves were almost superimposable; however, for 6mA- genes, the Δ*MLL1* curve was substantially higher than the WT curve downstream of TSS.

B. Differentially expressed genes in WT and Δ*MLL1*. Left: 6mA+ and 6mA- genes in WT. Right: H3K4me3+ and H3K4me3- genes in WT.

C. Relationship between changes in 6mA and gene expression in Δ*AMT6* (top) and Δ*AMT7* (bottom), normalized by WT. Δ*AMT6 or* Δ*AMT7* versus WT: $Log_2$(FoldChange) for individual genes in 6mA (x-axis: ΣP) and expression (y-axis: RPKM). Green: genes with significantly reduced 6mA (>256×) and increased expression levels (>2×). Red: genes with significantly reduced 6mA (>256×) and decreased expression levels (>2×).

**Table S1.** 6mA statistics.

| Single molecule | WT | | ΔAMT1 | | ΔAMT6 | | ΔAMT7 | |
|---|---|---|---|---|---|---|---|---|
| | Number | Percentage (%) | Number | Percentage (%) | Number | Percentage (%) | Number | Percentage (%) |
| **ApT** | 992,618,784 | 100.00 | 842,024,326 | 100.00 | 951,399,212 | 100.00 | 638,977,720 | 100.00 |
| **6mApT sites** | 18,750,787 | 1.89 | 4,248,723 | 0.50 | 16,020,425 | 1.68 | 3,386,207 | 0.53 |
| **Full** | 16,747,446 | 89.32 | 128,540 | 3.03 | 13,341,450 | 83.28 | 1,837,998 | 54.28 |
| **Hemi-W** | 998,137 | 5.32 | 2,051,797 | 48.29 | 1,335,265 | 8.33 | 773,635 | 22.85 |
| **Hemi-C** | 1,005,204 | 5.36 | 2,068,386 | 48.68 | 1,343,710 | 8.39 | 774,574 | 22.87 |
| **Total 6mApT** | 18,750,787 | 100.00 | 4,248,723 | 100.00 | 16,020,425 | 100.00 | 3,386,207 | 100.00 |

| Single molecule | ΔMLL1 | |
|---|---|---|
| | Number | Percentage (%) |
| **ApT** | 719,377,086 | 100.00 |
| **6mApT sites** | 15,513,221 | **2.16** |
| **Full** | 13,650,472 | 87.99 |
| **Hemi-W** | 919,330 | 5.93 |
| **Hemi-C** | 943,419 | 6.08 |
| **Total 6mApT** | 15,513,221 | 100.00 |

**Table S2**. UHPLC-QQQ-MS/MS analysis of 6mA level in overexpression of AMT6 and AMT7.

| Sample | 6mA (nmol/L) | dA (nmol/L) | 6mA/dA | Mean of 6mA/dA (%) |
|---|---|---|---|---|
| SB210 | 14.989 | 3254.353 | 0.004606 | 4.589 |
| | 18.34 | 4011.317 | 0.004572 | |
| *AMT7*-OE | 34.127 | 6601.992 | 0.005169 | 4.65 |
| | 17.944 | 3794.08 | 0.004729 | |
| | 1.817 | 448.366 | 0.004052 | |
| *AMT6*-OE | 19.171 | 4727.722 | 0.004055 | 3.87 |
| | 15.582 | 3954.169 | 0.003941 | |
| | 11.053 | 3048.919 | 0.003625 | |

**Table S3.** 6mA levels in 6mA+ and 6mA- genes, as well as rDNA.

| | | | 6mA+ | 6mA- | rDNA |
|---|---|---|---|---|---|
| **WT** | ΣP | | 524,769.10 | 9,262.63 | 2.55 |
| | Percentage (%) | Full | 80.74 | 41.38 | 38.68 |
| | | Hemi | 19.26 | 58.62 | 61.32 |
| ***ΔAMT1*** | ΣP | | 141,062.41 | 2,380.38 | 0.979 |
| | Percentage (%) | Full | 1.59 | 0.17 | 0.02 |
| | | Hemi | 98.41 | 99.83 | 99.98 |
| ***ΔAMT6*** | ΣP | | 471,436.88 | 5,019.98 | 1.36 |
| | Percentage (%) | Full | 72.13 | 26.02 | 6.49 |
| | | Hemi | 27.86 | 73.98 | 93.51 |
| ***ΔAMT7*** | ΣP | | 152,307.42 | 1,634.06 | 1.57 |
| | Percentage (%) | Full | 36.37 | 16.53 | 27.13 |
| | | Hemi | 63.63 | 83.47 | 72.87 |
| ***ΔMLL1*** | ΣP | | 554,765.50 | 31,624.65 | 2.48 |
| | Percentage (%) | Full | 78.01 | 58.55 | 21.71 |
| | | Hemi | 21.99 | 41.45 | 78.29 |
